# Disease-associated amino acid substitutions outside canonical motor motifs impair Myo1e localization and activity

**DOI:** 10.64898/2026.10.15.759198

**Authors:** Kazi Sayeeda, Jinghua Ge, Christopher M. Yengo, Mira Krendel

## Abstract

Steroid-resistant nephrotic syndrome (SRNS) is a severe kidney disorder characterized by proteinuria and disruption of glomerular filtration, often driven by genetic defects in genes encoding proteins required for podocyte functions. Among these, *MYO1E*, encoding the actin- and membrane-associated motor protein Myosin 1e (Myo1e), has been implicated in familial and sporadic forms of SRNS. Here, we investigated two *MYO1E* missense variants, A92E and G562R, identified in trans in a patient with childhood-onset SRNS and located outside canonical P-loop and switch subdomains of the Myo1e motor domain. Both residues are highly conserved, located within predicted α-helices in the motor domain, and replaced by larger charged residues, which may reduce protein stability or folding. In podocyte-derived cells, both mutants failed to localize at intercellular junctions and displayed reduced colocalization with actin. The G562R variant additionally formed cytoplasmic aggregates that were partially expression-level-dependent and exhibited decreased fluorescence recovery after photobleaching, indicating impaired protein dynamics. In vitro motility and ATPase assays indicated that the G562R mutant exhibited a marked loss of ATPase activity while the A92E mutant displayed ATPase activity similar to the WT but was characterized by reduced motility and thermal stability. Together, these results demonstrate that A92E and G562R mutations compromise Myo1e localization and function, supporting their pathogenic role in SRNS.

## INTRODUCTION

Nephrotic syndrome (NS) is a kidney disease characterized by proteinuria and hypoalbuminemia resulting from the loss of selectivity of the glomerular filtration barrier (GFB) (Preston et al., 2019). One of its histological subtypes, focal segmental glomerulosclerosis (FSGS), is characterized by glomerular scarring and progressive podocyte injury. Although no targeted therapies are currently available, NS is typically treated with corticosteroids and other immunosuppressants as first-line therapy (Bierzynska and Saleem, 2017; Varghese and Majumdar, 2022). Responsiveness to corticosteroid therapy serves as a key predictor of outcome: patients who respond rarely progress to kidney failure, whereas those with persistent proteinuria after 4–6 weeks of corticosteroid treatment, defined as steroid-resistant nephrotic syndrome (SRNS), often experience progressive renal decline despite alternative interventions such as renin–angiotensin system inhibitors or calcineurin inhibitors (Eddy and Symons, 2003; Ha, 2017; Trautmann et al., 2020). Most patients with SRNS ultimately progress to kidney failure within 5–15 years of diagnosis (Cason et al., 2023; Ha, 2017).

Genetic factors account for a major subset of SRNS cases, with 10-30% of non-familial SRNS cases being characterized by mutations in podocyte-associated genes (Ha, 2017; Trautmann et al., 2020). The most common histological correlate in these cases is FSGS. Thus, genetically determined NS is usually steroid-resistant, associated with FSGS, and often progresses to end-stage renal disease (ESRD) (Ha, 2017). SRNS with a genetic etiology is more prevalent in children than in adults, with many pediatric cases harboring genetic defects in podocyte-specific genes (Cason et al., 2023; Ha, 2017). Autosomal recessive SRNS typically presents in early childhood with high penetrance, whereas autosomal dominant forms often present later with incomplete penetrance, and individuals may remain asymptomatic (Preston et al., 2019).

Products of genes mutated in hereditary NS commonly regulate the podocyte actin cytoskeleton or slit diaphragm assembly (Hall and Gbadegesin, 2015). One such autosomal recessive gene, *MYO1E*, encodes Myosin 1e (Myo1e), a cytoskeletal scaffold and membrane-associated motor protein expressed predominantly in podocytes (Krendel et al., 2009; Mele et al., 2011; Sanna-Cherchi et al., 2011). As a class I myosin, Myo1e contains an N-terminal motor domain that binds actin filaments and hydrolyzes ATP, a neck domain that binds light chains and acts as a lever arm, and a long tail domain with TH1 (Tail Homology 1), TH2, and SH3 subdomains (McIntosh and Ostap, 2016) (**Figure 1A**). Through its tail domain, Myo1e binds negatively charged phospholipids, linking the plasma membrane to the actin cortex and supporting actin-activated ATPase activity that drives lamellipodia formation and cell migration (Noris and Remuzzi, 2012).

**Figure 1:**
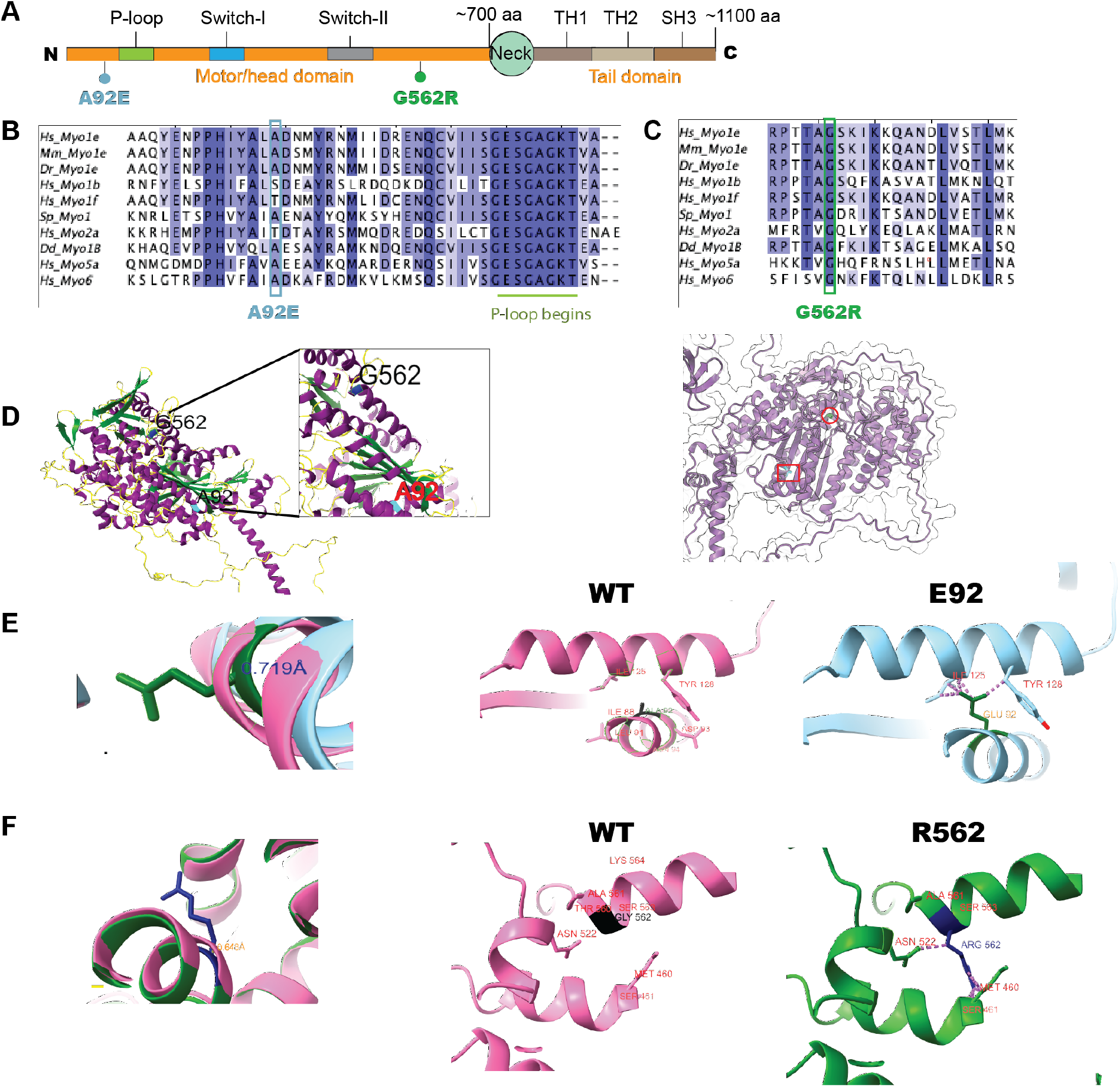
Structural context of SRNS-associated Myo1e variants. (**A**) **Domain map of human Myo1e**, indicating the locations of the SRNS-associated point mutations A92E and G562R. (**B and C**) **Multiple sequence alignments** of regions surrounding A92 (B) and G562 (C) sites across various myosins indicates that these amino acid positions are occupied by small amino acid residues across species and myosin isoforms. The blue box in (**B**) and the green box in (**C**) denote A92 and G562 residues, respectively. (**D**) **Secondary structure and solvent accessibility** of A92 and G562. Left: predicted structure of a portion of the Myo1e motor domain from AlphaFold2 where the secondary structure elements are color-coded, with the α-helices in purple, the β-sheets in dark green, and the loops in yellow. Both residues A92 (cyan) and G562 (dark blue) reside within predicted α-helices. Inset shows a zoomed-in view of the region containing the G562 and the A92 residues. Right: Solvent-accessibility analysis based on SASA (absolute solvent-accessible surface area) values determined using ChimeraX. The semi-transparent molecular surface (50% opacity) is overlaid on the Myo1e structure (shown in hot pink), with the positions of residues A92 (red square) and G562 (red circle) highlighted. The low SASA values for both residues indicate that A92 and G562 are buried within the protein core. (**E and F**) **Structural modeling** predicts that replacement of buried A92 or G562 residues with bulky charged residues introduces local steric clashes for A92E (**E**) and G562R (**F**) mutations. (**E**) Left: Structural overlay of wild-type (hot pink) and mutant Myo1e protein containing E92 (light blue), with side chains in dark green, showing local structural displacement near the mutation site, with a root-mean-squared deviation (RMSD) of 0.719 Å (dark blue). Middle: Wild-type Myo1e (hot pink) with A92 side chain in black, showing no steric clashes with neighboring residues. Right: Mutant Myo1e protein (light blue) with E92 side chain in dark green, forming interatomic clashes with I125 and Y128 (pink) with neighboring residues in red. (**F**) Left: Structural overlay of wild-type (hot pink) and mutant Myo1e protein containing R562 (dark green), with side chains in dark blue, showed local structural displacement near the mutation site, with RMSD of 0.648 Å (orange). Middle: Wild-type Myo1e (hot pink) with G562 side chain in black, showing no steric clashes with adjacent residues. Right: Mutant Myo1e protein (dark green) with R562 side chain in dark blue, introducing interatomic clashes with N522 and M460 (pink), and neighboring residues in red.

Myo1e dynamically localizes to lamellipodia tips and actin-rich puncta that precede adhesion maturation, implicating it in stabilizing matrix adhesions (Gupta et al., 2013). It associates with slit diaphragm proteins such as ZO-1, as well as with endocytic regulators including dynamin and synaptojanin-1 and with focal adhesion kinase (FAK) (Bi et al., 2013; Heim et al., 2017; Krendel et al., 2007), suggesting functions in cell–cell adhesion, membrane trafficking, and signaling. In the kidney, Myo1e is enriched at podocyte foot processes and junctions where it colocalizes with F-actin (Bi et al., 2013). Both complete and podocyte-targeted knockout of Myo1e results in proteinuria and podocyte dysfunction, including foot process effacement and glomerular basement membrane (GBM) thickening and splitting (Chase et al., 2012; Krendel et al., 2009). Myo1e-null mice develop proteinuria by three weeks of age (Krendel et al., 2009), and patients with *MYO1E* mutations also exhibit podocyte foot process effacement and characteristic GBM defects (Liu et al., 2024).

Previously analyzed *MYO1E* mutations, all identified in the homozygous state in the affected individuals, cluster in conserved subdomains of the motor domain (Krendel et al., 2023) such as the P-loop, switch-I, and switch-II regions, which mediate nucleotide binding, communication between the nucleotide and the actin-binding regions, and ATP hydrolysis (Diensthuber et al., 2024; Kintses et al., 2007; Mercier et al., 2015). Notable disease-associated mutations in *MYO1E* include T119I in the P-loop, A159P in Switch-I, and D388H in Switch-II (Krendel et al., 2023).

In contrast, the mutations analyzed in this study lie outside of these canonical motifs. Identified in a compound heterozygous state in a single patient, the A92E variant was inherited from the father and the G562R variant from the mother, both asymptomatic heterozygotes (Krendel et al., 2023). Despite not being associated with the well-characterized motor domain motifs, these mutations result in a histopathological phenotype similar to previously reported *MYO1*E mutations, including severe foot processes effacement, detachment, and irregular GBM thickening (Krendel et al., 2023). This study investigates how these newly identified *MYO1E* variants alter Myo1e structure, localization, and activity to provide functional evidence supporting their pathogenicity.

## MATERIALS AND METHODS

### PODOCYTE CULTURE

Conditionally immortalized podocyte-derived cells lacking Myosin1e (Myo1e-KO) were obtained from Myo1e-knockout (Myo1e-KO) mice (Krendel et al., 2009) crossed with Immortomouse (a transgenic mouse expressing a transgene encoding a thermosensitive, interferon-inducible variant of the SV-40-T-antigen), with isolation procedures originally described in (Bi et al., 2013). In brief, podocytes were isolated using the protocol described in (Mundel et al., 1997), involving sequential sieving of glomeruli. Individual podocyte clones were then obtained through dilution cloning. These clones were characterized and confirmed for Myo1e deficiency using immunostaining, immunoblotting, and RNA sequencing (Bi et al., 2013; Liu et al., 2022). While these cells expressed a number of markers characteristic of podocytes, they exhibited a gradual loss of the podocyte-specific marker nephrin, therefore, we designated them as podocyte-like cells (Liu et al., 2022). Cells were maintained in RPMI-1640 medium supplemented with 10% FBS, 1% antibiotic-antimycotic (ABM), and 50 units/mL IFN-γ (Cat # 11040596001; Batch # 68713300; Lot # 68713300, Millipore Sigma) at 33°C with 5% CO_2_ on 0.1 mg/mL collagen-I-coated (Rat tail Collagen Type 1; Cat # 354236, Lot # 17323009, Corning) 60 mm tissue culture dish (Cat # 430166, Corning). Cells were subcultured every four days and maintained below 80% confluency.

For differentiation, cells were maintained at 37°C in the absence of IFN-γ. At differentiation day 10, cells were infected with adenoviruses to express recombinant proteins for live-cell imaging. Following imaging, cells were collected for protein analysis via a Western blot.

### CONSTRUCTS

Adenoviral constructs for EGFP-Myo1e expression (WT and mutants) were generated by VectorBuilder using an adenoviral backbone containing the human cytomegalovirus (CMV) promoter for strong expression and an N-terminal EGFP tag. An ampicillin-resistance marker enabled bacterial selection for DNA amplification. Verified constructs were packaged by Vectorbuilder and shipped ready for in vitro transduction. The mCherry-LifeAct adenoviral construct was previously described (Bi et al., 2014).

For baculoviral expression, mutant Myo1e fragments were cloned into the pFastBac vector containing C-terminal EGFP-FLAG-AviTag sequences (Liu et al., 2022). Mutations were introduced using the Q5 High-Fidelity Site-Directed Mutagenesis Kit (NEB M0493). All plasmid constructs were verified by whole-plasmid sequencing (Plasmidsaurus). For bacmid-producing recombination, the validated plasmid was introduced into MAX Efficiency DH10bac^TM^ competent cells (Lot# 2732903, Gibco). Successful recombination was confirmed by blue-white colony selection on plates containing kanamycin, gentamycin, and tetracycline. A white colony was selected, and recombined baculoviral DNA was extracted using the PureLink^TM^ HiPure Plasmid Miniprep Kit (Ref# K2100-02, Lot# 2787626).

To further confirm the successful recombination, PCR amplification of the insert was performed, and the fragment size was assessed via agarose gel electrophoresis. Once the correct band was identified, it was purified using the NucleoSpin Gel and PCR Clean-up kit ( Macherey-Nagel, Ref# 740609.50 Lot# 1704/002, Takara Bio USA, Inc.), followed by sequencing through Plasmidsaurus. Primers used for mutagenesis are listed in Supplemental Table 1.

### INFECTING DIFFERENTIATED PODOCYTES WITH ADENOVIRAL VECTORS FOR LIVE-CELL IMAGING

A total of 1.2×10^4^ undifferentiated podocyte-derived cells were seeded in a 35 mm glass-bottom dish (P35G-1.5-14-C, Mattek Corporation) coated with 10 μg/mL collagen-IV (Cat # 354233, Lot # 7114009, Corning mouse collagen) and maintained at 37°C without IFN-γ. On the 10^th^ day of differentiation, adenoviral transduction was performed following the protocol described in (Bi et al., 2014) to express fluorescently-tagged proteins for live-cell imaging. At the time of infection, podocyte-derived cells were approximately 70-80% confluent, and imaging was conducted 24 hours post-infection.

For infection, the virus was diluted in complete medium (RPMI with 10% FBS and 1% ABM) at a concentration of 1×10^6^ PFU/mL for live-cell imaging. For experiments using reduced Myo1e^G562R^ expression, the G562R virus was further diluted twofold (1:200), corresponding to a final concentration of 5 x 10^5^ PFU/mL. When expressing multiple markers, two viruses were combined and used simultaneously. The culture medium was removed and replaced with the diluted virus in a minimal volume of 200 μL, ensuring full coverage of the cell monolayer. Cells were incubated at 37°C for 1 hour, with gentle rocking every 20 minutes. 300 μL of medium was added every 20 minutes to prevent drying out due to low volume. After 1 hour of incubation, 1.5 mL of complete medium was added, and the cells were kept overnight.

After 24 hours, before imaging, the medium containing the virus was carefully removed to prevent spillage. The cells were then washed three times with 1 mL of complete medium. Following the washes, 2 ml of fresh complete medium was added, and the cells were prepared for imaging.

### CONFOCAL MICROSCOPY FOR JUNCTIONAL ENRICHMENT ANALYSIS

For the analysis of cellular junctional enrichment and fluorescence recovery after photobleaching (see next section), imaging was performed using a PerkinElmer UltraView VoX Spinning Disk Confocal Microscope mounted on a Nikon Eclipse Ti-E microscope equipped with a Hamamatsu C9100-50 EMCCD camera and a Nikon Apo TIRF 60x (1.49 N.A.) oil objective, controlled by Volocity software. Podocyte-derived cells were imaged at 60X magnification in complete RPMI at 36°C.

Laser power and exposure time were optimized using cells expressing WT Myo1e and kept constant for all samples. Dual-channel fluorescence imaging was performed with the following acquisition settings: 561 nm (red channel, mCherry-LifeAct): 50% laser power, 500 ms exposure; 488 nm laser (green channel, EGFP-Myo1e): 34.5% laser power, 300 ms exposure. With these settings, the maximum fluorescence intensities were 6000 for the red channel, 7700 for the green channel, and 1800 for cytoplasmic green signal. These values were used as a reference point to avoid oversaturation across all samples.

Z-stacks spanning 4 µm (0.3 µm spacing between images) were acquired from the mid-cell plane. Images were analyzed in Fiji using a mid-plane section. A rectangular ROI was drawn around cell-cell junctions in the red (actin) channel, then applied to the green (Myo1e) channel, and additional ROIs were selected for cytosol and background regions. Junctional enrichment was calculated as the ratio of background-corrected junctional to cytosolic fluorescence. Data from three independent experiments with six cells per experiment were analyzed using one-way ANOVA versus WT control.

Line-scan analysis was used to examine Myo1e colocalization with actin. In Fiji, a 5 µm × 0.01 µm line was drawn perpendicular to the junction in a mid-cell confocal section. Fluorescence profiles were extracted and normalized to each cell’s maximum intensity (0–1 range). For consistency, junctions oriented horizontally were rotated 90° prior to analysis without altering content. Fluorescence values from 18 cells per mutant construct (A92E and G562R) and 21 cells expressing WT were plotted in GraphPad Prism as XY data (distance vs. intensity). Mean ± SD were calculated and displayed with error bars.

### FLUORESCENCE RECOVERY AFTER PHOTOBLEACHING AND ANALYSIS

Photobleaching was performed with a 488 nm laser targeting 50 × 50-pixel ROIs within cytoplasmic aggregates. Cells were photobleached for 2 seconds, and fluorescence recovery images were acquired at 0.5 s intervals for 5.5 s, followed by imaging at 1 s intervals for an additional 60 seconds.

Integrated density (area x mean fluorescence intensity) was measured for bleached and control unbleached ROIs throughout the experiment, and background correction was applied. The background-corrected integrated density of the control ROI for each timepoint was used to correct for bleaching during imaging/recovery. Recovery was normalized to the mean of the two pre-bleach values.

Normalized postbleach recovery curves from individual cells (three cells per construct, images collected on the same day) were fitted separately in GraphPad Prism 9.0 using a one-phase association model: Y= Y_0_ + (Plateau - Y_0_) * (1-exp(-K*X)) with Y_0_ constrained to 0, where the plateau is the Y value at infinite times expressed in the same unit as Y. To compare the construct of interest with the Myo1e-WT, differences in plateau values (mobile fraction) were examined. Because these measurements were obtained from one experiment, comparisons among conditions were considered descriptive.

### TOTAL PROTEIN STAINING AND WESTERN BLOT ANALYSIS

After imaging, cells were lysed in cold GFP-Trap Lysis Buffer (10 mM Tris-HCl [pH 7.5], 150 mM sodium chloride, 0.5 mM EDTA, 0.5% Nonidet P-40 and 0.09% sodium azide) with protease inhibitor (Pierce Protease inhibitor Cat No. A32965, Thermo Scientific). After addition of the lysis buffer, the cells were lifted using a cell scraper and boiled at 100°C for 5 minutes after the addition of the 5x sample buffer.

Proteins were separated by SDS-PAGE (170 V, 1 h 17 min, in 15% acrylamide gel) and transferred to PVDF membranes. Equal protein loading was confirmed by SYPRO Ruby Protein Blot stain (Lonza #50565, Lot #2921590) per manufacturer’s protocol. Membranes were air-dried and fixed (7% acetic acid, 10% methanol, 15 min), washed, stained with SYPRO Ruby Protein Blot stain for 15 min, and imaged on an Azure 600 Imager.

Membranes were then blocked with 5% milk in TBST and probed with rabbit anti-GFP (Thermo A6455; RRID: AB_221570) followed by goat anti-rabbit HRP (Jackson ImmunoResearch) and WesternBright ECL substrate (Advansta K-12045-D20). Chemiluminescent intensities were normalized to total protein using Image Lab software (WT set to 1).

### BACULOVIRUS EXPRESSION AND PURIFICATION OF MYO1E CONSTRUCTS

Recombinant human Myo1e, truncated after the IQ domain (residues 1-725), with C-terminal GFP, Avi, and FLAG (DYKDDDDK) tags, was expressed in Sf9 insect cells using the Bac-to-Bac baculovirus expression system (Liu et al., 2022; Liu et al., 2024; Sun et al., 2006; Trivedi et al., 2013). All Myo1e constructs were co-expressed with calmodulin to ensure proper folding and stability of the IQ motif, as it is standard in myosin expression systems (El Mezgueldi et al., 2002).

For protein production, Sf9 cultures were infected with P1 viral stocks and incubated at 27°C for 72 hrs. Cells were harvested by centrifugation, and protein lysates were subjected to affinity purification using anti-FLAG M2 resin (Lot No.#0000378819, Sigma Aldrich), followed by elution with 3X FLAG peptide (El Mezgueldi et al., 2002; Liu et al., 2024; Sun et al., 2006; Trivedi et al., 2013).

The purity was greater than 95% based on Coomassie-stained SDS gel. Myo1e concentrations were determined using a Bio-Rad microplate assay using bovine serum albumin as a standard (Sun et al., 2006; Yengo et al., 2012). Actin was purified from rabbit skeletal muscle using an acetone powder method as described (Pardee and Spudich, 1982).

### *IN VITRO* MOTILITY ASSAY

The *in vitro* motility assay was used to assess the actin filament gliding activity of MYO1E-motor-IQ-WT-EGFP-FLAG-Avi, A92E, and G562R constructs, as described previously (Dantas et al., 2018; Raval et al., 2016; Yengo et al., 2012). Following the approach established in our lab (Liu et al., 2022), Myo1e constructs were adhered to nitrocellulose-coated coverslips via an anti-GFP antibody (0.1 mg/mL; Invitrogen^TM^, G10362), which provides strong and reliable attachment without requiring additional biotin-streptavidin interactions. The coverslips were then blocked with 2 mg/mL bovine serum albumin (BSA) prepared in KMg50 buffer containing 2 mM MgCl_2_ (50 mM KCl, 1 mM EGTA, 2 mM MgCl_2_, 1 mM dithiothreitol, 10 mM imidazole [pH 7.0]).

To block inactive or non-functional myosin heads, 2 μM sheared, unlabeled F-actin was added, followed by incubation with 2 mM ATP to help release active heads while inactive heads remained bound to actin in the presence of ATP. Next, phalloidin-labeled F-actin filaments were introduced into the flow chamber. Immediately prior to imaging, an activation buffer was added. This motility buffer consisted of KMg50 containing 2 mM MgCl_2_ supplemented with 2.5 μM calmodulin, 0.35% methylcellulose, 0.45 mM phosphoenolpyruvate, 45 U/mL pyruvate kinase, 0.1 mg/mL glucose oxidase, 1 mg/mL glucose, 0.018 mg/mL catalase, and 2 mM ATP to support ATP regeneration and minimize photobleaching during video acquisition (Yengo et al., 2012).

Actin filament motility was visualized using a Leica DMi8 fluorescence microscope equipped with a 63x/1.4 N.A. phase-contrast objective, a DsRed filter, and a Hamamatsu ORCA Flash4.0 CMOS camera. Time-lapse images were acquired at 10-second intervals for up to 10 minutes. Gliding velocities were determined by manually tracking individual actin filaments using the MTrackJ plugin in ImageJ (Meijering et al., 2012).

### STEADY-STATE ATPASE MEASUREMENTS

The actin-activated ATPase activities of purified Myo1e motor-IQ domain constructs (WT, A92E, and G562R) were measured using an NADH-coupled enzymatic assay, as previously described (De La Cruz et al., 2000; Tang et al., 2021). All reactions were performed at 25°C in KMg50 buffer. This assay couples ATP hydrolysis to the oxidation of NADH through an ATP regeneration system, allowing ATPase activity to be monitored by the decrease in absorbance at 340 nm.

Purified Myo1e (0.1 µM final concentration) was premixed with varying concentrations of F-actin (0-60 µM) and an NADH-coupled assay mixture containing NADH, phosphoenolpyruvate (PEP), lactate dehydrogenase, and pyruvate kinase in KMg50 buffer. Reactions were initiated by adding ATP immediately before data acquisition. Control reactions lacking Myo1e were used to subtract background NADH oxidation, which was consistently <1%.

Absorbance changes were recorded over a 200 s time course using an Applied Photophysics stopped-flow apparatus, with data collected every 0.2 s. A standard curve of known ADP concentrations was used to convert absorbance units to molar ATP turnover. ATPase rates were plotted against actin concentration and fitted to the Michaelis-Menten equation (V=V_0_+k_cat_*[actin]/(K_ATPase_+[actin]), to determine the maximal actin-activated ATPase rate (k_cat_), and half-maximal (K_ATPase_) actin concentration. Basal ATPase rates were measured using 0.5 µM myosin in the absence of actin. The total theoretical maximal ATPase activity (V_max_) was calculated as the sum of basal and actin-activated activities (V_max_ = V_0_ + k_cat_). Additionally, experimental ATPase rates at a saturating actin concentration of 60 µM were reported for direct comparison. All experiments were repeated with at least three independent protein preparations, and statistical comparisons between WT and mutants were made using unpaired t-tests.

### PROTEIN THERMAL SHIFT ASSAY

Thermal stability of purified Myo1E WT, A92E, and G562R constructs was assessed using a fluorescence-based protein thermal shift assay. Each protein was diluted to a final concentration of 1 µM in KMg150 buffer containing 2 mM ATP and mixed with SYPRO Orange at a final 2× concentration. Samples were transferred to a qPCR-compatible plate and analyzed using a QuantStudio™ 3 Real-Time PCR System. The temperature was increased from 25°C to 95°C at a rate of 0.5°C/min, with fluorescence monitored throughout the temperature ramp. For each sample, fluorescence values were normalized between the minimum and maximum signals. The normalized unfolding curves were fitted using a Boltzmann sigmoidal equation, and the melting temperature (Tm) was defined as the midpoint of the fitted transition. Three technical replicates were analyzed for each construct, and Tm values are reported as mean ± SD.

### BIOINFORMATIC PREDICTION OF MUTATIONAL EFFECTS

Multiple protein sequence alignments were performed using the MUSCLE (Multiple Sequence Comparison by Log-Expectation) algorithm within MacVector (version 17.0.10). The NCBI reference sequence accession numbers of all aligned protein sequences are listed in Supplemental Table 2.

To investigate the structural context of residues A92 and G562 in human Myo1e, we used AlphaFold2 via the Neurosnap web server, which utilizes ColabFold for accelerated and accurate structure prediction (Mirdita et al., 2022). The input was the full-length FASTA sequence of human Myo1e (UniProt ID: Q12965). Among the models generated for each construct, the top-ranked models were selected based on the mean predicted Local Distance Difference Test (pLDDT) scores. The selected models each had mean pLDDT scores >80, representing high model confidence (Mirdita et al., 2022).

A92E and G562R substitutions were introduced into the FASTA sequence for qualitative visualization of the local side-chain environment using Benchling. The modified sequences were then re-submitted to the same AlphaFold2 pipeline to obtain mutant structural models. The resulting PDB models (Mirdita et al., 2022), both wild-type and mutant, were imported into UCSF ChimeraX (version 1.10.1) for all downstream structural analysis.

Within ChimeraX, we assessed the type of secondary structure each residue lies within and evaluated whether the residues were surface-exposed or buried by calculating solvent-accessible surface area (SASA) per residue (Ali et al., 2014).

SASA was computed using the rolling-probe method (Shrake-Rupley algorithm) with a 1.4 Å probe radius (Tien et al., 2013), which approximates the size of a water molecule. Calculations were performed over the entire folded protein structure, restricted to protein atoms, to obtain accurate per-residue SASA values that reflect each residue’s contribution to the overall area of the protein (Meng et al., 2023; Pettersen et al., 2021). We intentionally avoided isolated, residue-only SASA calculations, as these tend to overestimate exposure by ignoring the spatial context of the folded structure (Meng et al., 2023). To further interpret the surface accessibility, we normalized the per-residue SASA values against known theoretical maximum ASA values to compute relative solvent accessibility (RSA) (Savojardo et al., 2020; Tien et al., 2013). Based on standard thresholds, residues were classified as buried if RSA<0.20 (20%) and exposed if RSA ≥ 0.20 (Savojardo et al., 2020).

Finally, to assess the structural consequences of the mutations, we aligned the wild-type and mutant structures to evaluate local structural displacement and identified steric clashes introduced in the immediate environment of the mutated residues (Xiong et al., 2022).

### STATISTICAL ANALYSIS

All statistical analyses and graphing were performed using GraphPad Prism 9.0 software. A one-way ANOVA with Dunnett’s multiple comparisons test was used to analyze data for multiple comparisons between WT and the mutants in podocyte experiments. For Western blot data analysis, we used a two-tailed one-sample t-test. ATPase, motility, and thermal stability values for the WT vs. mutant constructs were compared using unpaired t-tests. Statistical significance was set at *P≤* 0.05 for all analyses.

## RESULTS

### *MYO1E* VARIANTS

A previous clinical case series identified a patient carrying two heterozygous missense variants in the *MYO1E* gene, A92E inherited from the father and G562R from the mother (Krendel et al., 2023). As shown in **Figure 1B,C**, multiple sequence alignment across human Class I myosins, other non-muscle myosins, and Myo1e orthologs from various species demonstrated that the A92 amino acid residue is evolutionarily conserved within Myo1e but not across all Class I myosins or other non-muscle myosins, whereas the G562 amino acid residue is conserved across Myo1e orthologs and in other non-muscle myosins. Notably, the position corresponding to alanine 92 in Myo1e (A92) is occupied by alanine, serine, or threonine in other myosins examined, consistent with the conservation of a relatively small, uncharged residue at this site. Similarly, the position corresponding to G562 in Myo1e is most often occupied by glycine, or in a few cases (not shown in the figure) alanine, indicating strong conservation of a small residue. Together, these observations suggest that substitutions introducing larger or charged residues at these positions may disrupt normal motor domain structure or function.

Mapping the A92 and G562 residues onto an AlphaFold2 model of full-length human Myo1e placed both within predicted α-helical regions of the motor domain (**Figure 1D**). Both residues were predicted to be buried in the WT model, with relative solvent accessibility (RSA) values of close to zero (0.00 for A92 and 0.01 for G562). RSA values range from 0 (fully buried) to 1 (fully exposed, 100%) (Gress and Kalinina, 2020; Xiong et al., 2022) (Savojardo et al., 2020). Replacing these residues with the larger, charged amino acids glutamate or arginine could therefore destabilize local packing or helical stability due to steric hindrance or electrostatic repulsion. The positions corresponding to Myo1e A92 and G562 are occupied by a threonine (T154) and a glycine (G647), respectively, in human nonmuscle myosin-2A encoded by the MYH9 gene, and the experimentally determined structure of non-muscle myosin-2A (Heiringhoff et al., 2025) also places these homologous residues in buried α-helices. A similar mutation of the homologous buried residue in Myo5b (A143E) has been linked to microvillus inclusion disease (van der Velde et al., 2013), further highlighting the importance of a small uncharged residue in this position.

The position of amino acid residues relative to the protein core or surface is an important determinant of their functional role, evolutionary constraint, and susceptibility to pathogenic mutations (Aloy et al., 2001; Gress and Kalinina, 2020). While mutations of residues on the surface can impact the specificity of protein interactions, mutations of residues in the protein core can influence overall stability and, in extreme cases, lead to misfolding and aggregation (Gress and Kalinina, 2020; Xiong et al., 2022). In line with this, the buried nature of A92 and G562, combined with their location in structured alpha-helical elements, suggests that their substitution could introduce strain or destabilize intramolecular contacts critical for Myo1e function.

Qualitative side-chain placement in ChimeraX identified potential steric incompatibilities between E92 and nearby I125/Y128 and between R562 and nearby N522/M460 (**Figures 1E** and **1F**). These findings support a model in which missense variants compromise Myo1e stability through local structural distortion and packing defects.

### EXPRESSION OF EGFP-TAGGED MYO1E ADENOVIRAL CONSTRUCTS

Adenoviral constructs encoding EGFP-tagged Myo1e^WT^ and mutant variants were transduced into Myo1e-KO podocyte-derived cells for localization assays (**Figure 2**). Western blot analysis (**Figure 2A**) confirmed the presence of full-length EGFP-Myo1e fusion proteins in Myo1e^WT^, Myo1e^A92E^, and Myo1e^G562R^-transduced cells. Although faint bands corresponding to EGFP and degraded EGFP-Myo1e products were observed, the most intense bands aligned with the 150 kDa molecular weight of the full-length fusion protein when probed with an anti-GFP antibody. When normalized to total protein loading (detected using SYPRO Ruby staining), quantitative analysis showed that Myo1e^A92E^ expression was not significantly different from Myo1e^WT^ (**Figure 2B**), whereas Myo1e^G562R^ demonstrated significantly higher expression under the transduction conditions used (**Figure 2C**). This elevated expression of Myo1e^G562R^ may contribute to the formation of cytoplasmic aggregates (**Figure 2D**).

**Figure 2:**
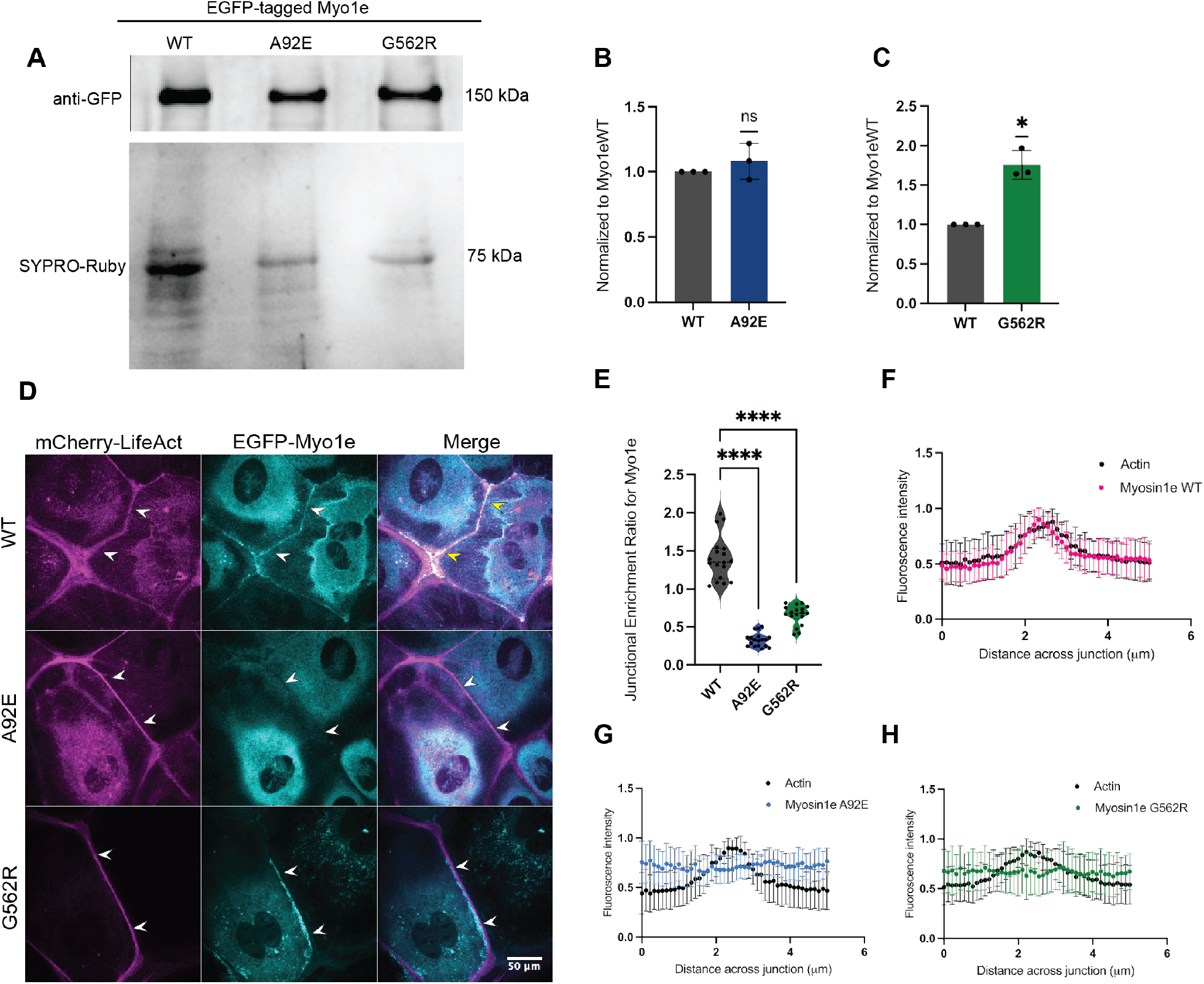
Expression and localization of WT and variant Myo1e in Myo1e-KO podocyte-derived cells. **(A) Western blot analysis of Myo1e expression.** Total cell lysates from podocytes expressing EGFP-Myo1e^WT^, A92E, or G562R were probed with anti-GFP antibody. SYPRO-Ruby staining was used for total protein normalization. **(B- C) Quantification of protein expression.** Densitometric analysis of Western blot bands from **(A),** normalized to Myo1e^WT^ levels (mean±SD from three independent experiments). Myo1e^A92E^ shows no significant change (p=0.4201) **(B),** while Myo1e^G562R^ is expressed at significantly higher levels (p=0.0191) (*p<0.05) **(C).** (**D) Confocal imaging of Myo1e localization.** Single plane confocal images of Myo1e-KO podocytes co-expressing mCherry-LifeAct (magenta) and EGFP-tagged Myo1e variants (cyan): wild-type (WT), A92E, or G562R. White arrowheads mark cell-cell junctions; yellow arrowheads indicate colocalization of Myo1e with actin. Scale bar, 50 µm. **(E) Quantification of junctional enrichment.** Violin plots show the distribution of the fluorescence intensity ratio of junctional to cytoplasmic Myo1e measured in individual cells (three independent experiments, 6 cells per experiment). Myo1e^WT^ is significantly enriched at the junctions compared to both Myo1e^A92E^ and Myo1e^G562R^ (****p<0.0001). **(F-H) Line scan analysis of junctional colocalization** shows that WT Myo1e (**magenta, F**) displays a distinct fluorescence intensity peak overlapping with actin (**black**), indicating strong colocalization at the junction. In contrast, A92E (**blue, G**) and G562R (**green, H**) display flat intensity profiles with no distinct peaks and the absence of elevated intensity at the peak of F-actin fluorescence (three independent experiment, 18 cells per mutant construct (A92E and G562R) and 21 cells expressing WT with error bars showing mean ± SD).

### MIS-LOCALIZATION OF MYO1E VARIANTS FROM CELLULAR JUNCTIONS

In glomeruli, Myo1e is associated with the slit diaphragm fraction and localizes along the foot process plasma membrane adjacent to slit diaphragms (Bi et al., 2013). In cultured podocyte-derived cells, Myo1e is enriched at the cell-cell junctions (Bi et al., 2013; Mele et al., 2011). Previous studies investigating Myo1e mutations found in pediatric FSGS patients revealed that these variants were primarily mis-localized to the cytoplasm, away from the cellular junctions. Since Myo1e plays a crucial role in actin reorganization during contact formation in podocytes, its mislocalization could contribute to disease pathology (Bi et al., 2013).

However, not all disease-associated Myo1e mutants are completely mis-localized. For instance, Myo1eD388H, described in (Liu et al., 2022), was found at cellular junctions colocalizing with actin filaments, while exhibiting loss of motor activity in vitro. This suggests that some pathogenic mutants may still localize correctly but could be functionally impaired in other ways. Therefore, it is essential to assess both localization and functional activity to understand their role in disease mechanisms.

To identify cell contacts, we used mCherry-LifeAct to label actin filaments, which are enriched at cell-cell junctions. Our analysis showed that both mutants, Myo1e^A92E^ and Myo1e^G562R^, failed to localize at the junctions (**Figure 2D, E**), instead being predominantly localized in the cytoplasm. To further examine Myo1e localization, we performed a line-scan analysis of EGFP-Myo1e intensity across cell junctions. The Myo1e^WT^ colocalized with F-actin at cell-cell contacts and exhibited a peak of intensity at the junction (**Figure 2F**). In contrast, the two mutants, Myo1e^A92E^ and Myo1e^G562R^, exhibited a much flatter intensity profile lacking a peak of brighter fluorescence coinciding with the peak of F-actin fluorescence at the junctions or in adjacent regions (**Figure 2G, H**). Unlike Myo1e^WT^, the mutants did not display strong peaks at these sites, (**Figure 2F-H**), indicating that mutant Myo1e constructs are more diffusely distributed in the cytoplasm rather than being localized at junctions. Although some Myo1e^G562R^-positive aggregates appeared to overlap with cell-cell junctions in the images, closer inspection showed that they were positioned adjacent to, rather than within, the F-actin-defined junction.

### EXPRESSION-DEPENDENT AGGREGATION OF Myo1e^G562R^

Myo1e^G562R^ but not Myo1e^A92E^ or Myo1e^WT^, formed cytoplasmic puncta under the initial transduction condition and was more highly expressed (Figure 2C). To determine whether this aggregation reflected an intrinsic property of the mutant protein or resulted from overexpression, we transduced cells with a lower titer of Myo1e^G562R^ virus and examined protein localization (**Figure 3A**). Reducing the amount of virus used decreased the extent of visible aggregates but did not restore proper junctional localization (**Figure 3B, C**), suggesting that while overexpression contributes to aggregation it does not account for the underlying mislocalization defect. Western blot analysis of EGFP-Myo1e^G562R^ showed that cells transduced at the lower viral titer expressed the full-length protein at levels comparable to EGFP-Myo1e^WT^ (**Figure 3D, E**).

**Figure 3:**
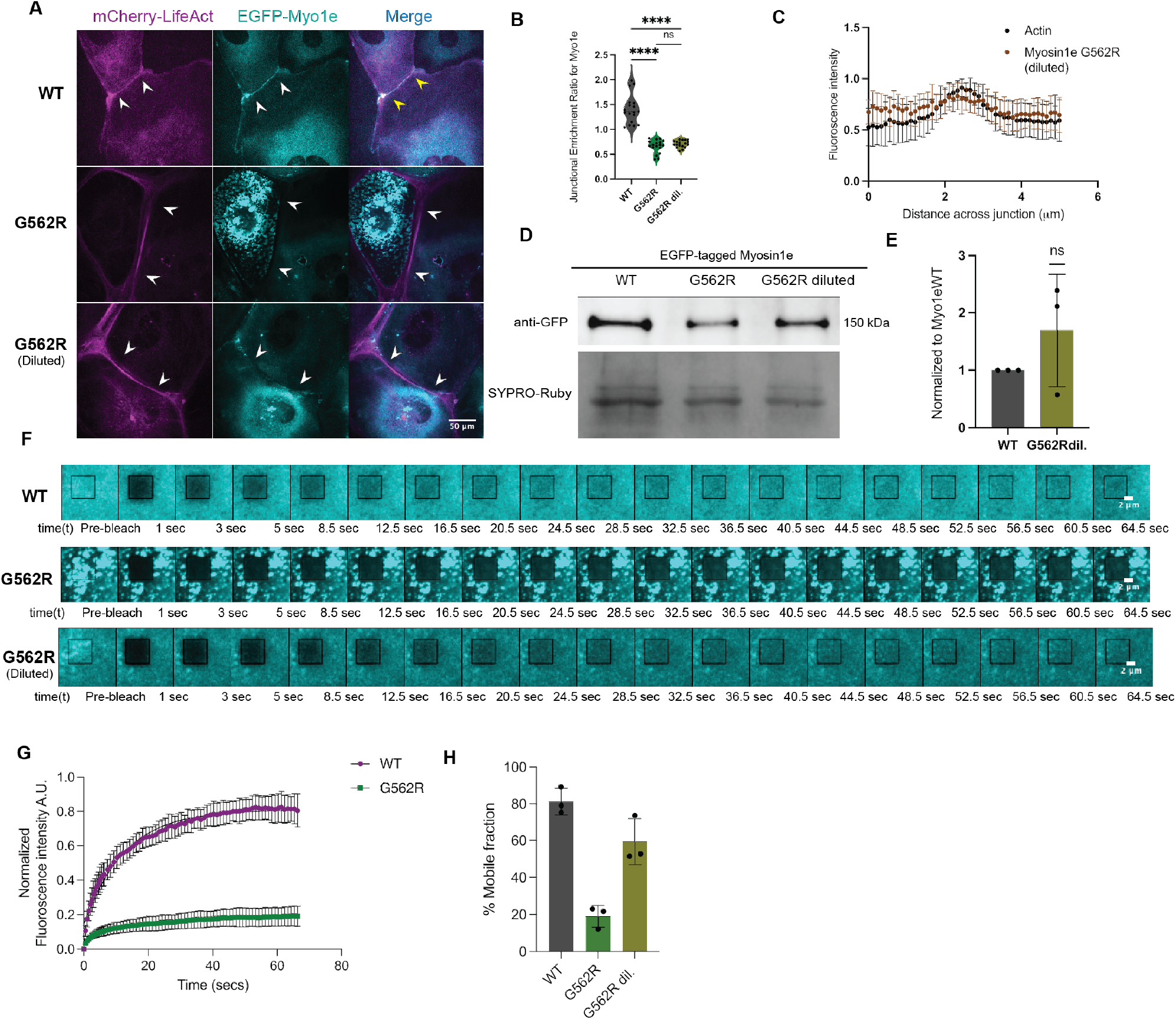
Concentration-dependent aggregation of Myo1e^G562R^. (**A**) **Confocal imaging of Myo1e Localization.** Single-plane confocal images of Myo1e-KO podocytes co-expressing mCherry-LifeAct (magenta) and EGFP-Myo1e variants (cyan): wild-type (WT), G562R, G562R (diluted). White arrowheads indicate cell junctions; yellow arrowheads indicate colocalization of actin and Myo1e. While EGFP-Myo1e^WT^ is enriched at the plasma membrane and strongly colocalizes with actin, the G562R mutant shows diffuse cytoplasmic localization with prominent aggregate-like structures, some of which are located along the cell-cell junction. Diluting the virus encoding the mutant construct reduces aggregate formation but does not restore junctional enrichment. Scale bar, 50 µm. (**B**) **Quantification of junctional enrichment**. Violin plot showing the ratio of Myo1e fluorescence intensity at cell-cell junctions relative to cytoplasmic regions (n=18 cells per condition, from three independent experiments; data points used for the WT and G562R are the same as in Figure 2) (****p<0.0001). (**C**) **Line-scan analysis of junctional colocalization.** Quantification of actin and Myo1e fluorescence intensity profiles along the junctions shown in (**A**), confirming decreased colocalization in G562R (diluted)-expressing cells. (**D**) **Western blot analysis of Myo1e expression.** Cell lysates were probed with anti-GFP antibody, loading was normalized using total protein stain. (**E**) **Quantification of Myo1e protein levels.** Quantification of band intensity in (**D**), normalized to the Myo1e^WT^ levels (mean±SD). Expression levels of the EGFP-Myo1e^G562R^ diluted construct are comparable to those of the EGFP-Myo1e^WT^ (p=0.3448). (**F**) **FRAP assay of cytoplasmic Myo1e dynamics.** FRAP images showing recovery of EGFP-Myo1e signal in cytoplasmic regions. The black rectangle indicates the bleached region. Scale bar, 2 µm. (**G**) **Fluorescence recovery curves.** Time course of fluorescence recovery for cytoplasmic Myo1e WT and G562R following FRAP. The data represent three cells/ROIs per condition (n=3, from one experiment, summary curves with error bars representing mean ±SD). (**H**) **Mobile fraction of cytoplasmic Myo1e.** The bar graph shows the recovering (mobile) fraction of Myo1e protein as determined by FRAP analysis (n=3 cells per condition, from a single experiment. Bars represent mean ±SD).

To clarify the nature of the observed punctate aggregates of the EGFP-Myo1e^G562R^, we performed FRAP analysis. FRAP measurements were performed on Myo1e^G562R^-positive puncta in the cytosol and on the cytoplasmic EGFP-Myo1e^WT^. In cells expressing Myo1e^WT^, cytoplasmic EGFP fluorescence recovered to approximately 80% after photobleaching. In contrast, Myo1e^G562R^ mutant displayed much lower recovery to approximately 20% of the prebleach fluorescence level (Figure 3F,G). Upon reducing expression levels by virus dilution, Myo1eG562R showed improved fluorescence recovery of ∼60% (Figure 3H), consistent with reduced aggregation. This suggests that cytoplasmic aggregation was partially driven by overexpression.

### MUTATIONS AFFECT MYO1E MOTOR ACTIVITY

Biochemical characterization of the Myo1e motor-IQ recombinant proteins purified from insect cells revealed distinct effects of the two variants on the steady-state ATPase activity (Figure 4, Table 1). Myo1e^WT^ had a basal (in the absence of actin) ATPase rate of 0.85 ± 0.07 s⁻¹ and its ATPase activity was further increased in the presence of actin. Myo1e^A92E^ retained a similar basal ATPase rate (0.92 ± 0.08 s⁻¹) but its actin-dependent activation was slightly less pronounced; nevertheless, its measured rate at 60 µM actin (1.05 ± 0.16 s⁻¹) was close to the WT (1.15 ± 0.13 s⁻¹). By contrast, Myo1e^G562R^ showed very low basal and actin-activated ATPase rates, with the rate of 0.12 ± 0.01 s⁻¹ at 60 µM actin.

**Figure 4.**
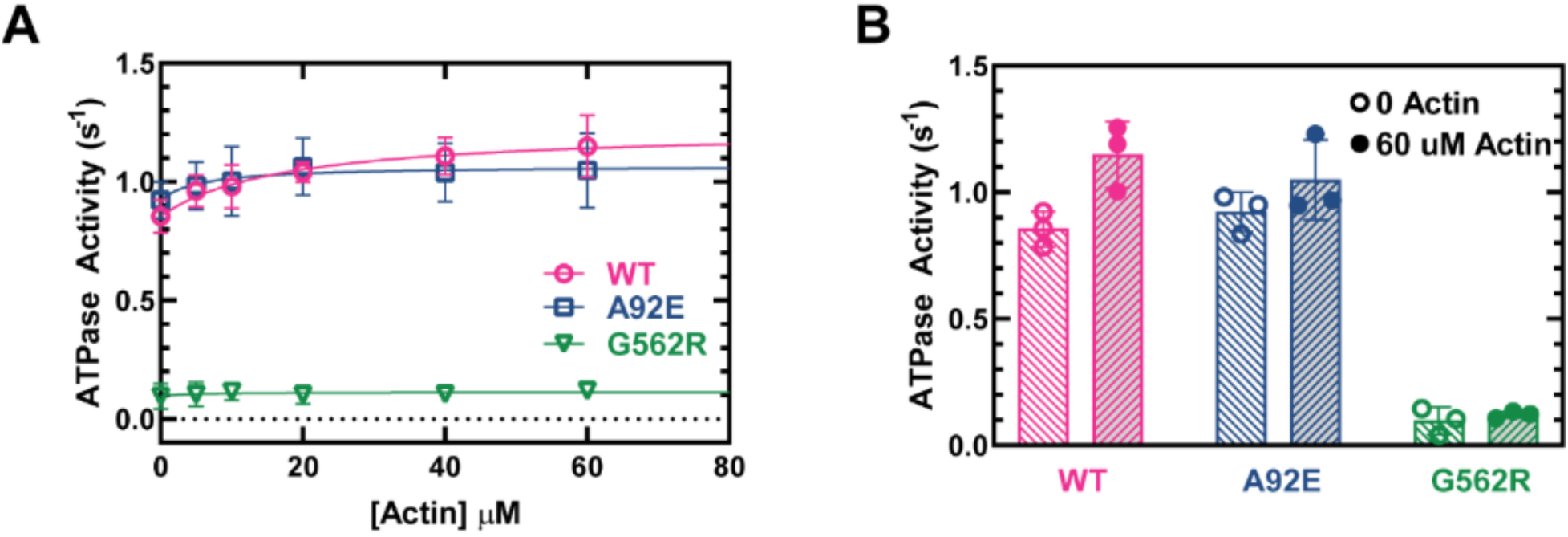
Actin-activated ATPase activity of Myo1e WT and disease-associated variants. (A) Steady-state ATPase activity of purified Myo1E WT, A92E, and G562R constructs measured at actin concentrations ranging from 0 to 60 µM. Data were fitted using the Michaelis–Menten equation. (B) Comparison of ATPase activities measured in the absence of actin (basal activity) and in the presence of 60 µM actin. Individual measurements are shown as open circles (0 µM actin) or filled circles (60 µM actin), and bars represent the mean ± SD. WT, A92E, and G562R shown in magenta, blue, and green, respectively. Data were obtained from three independent protein preparations (n = 3).

**Table 1.** Summary of basal and actin-activated ATPase activities of Myo1E^WT^ and disease-associated variants.

| Construct | $v_0$ ($\text{s}^{-1}$) | $k_{\text{cat}}$ ($\text{s}^{-1}$) | $v_{\text{max}}=v_0+k_{\text{cat}}$<br>($\text{s}^{-1}$) | $v_{\text{max}}$ ($\text{s}^{-1}$)<br>@60 $\mu\text{M}$ actin | $K_{\text{ATPase}}(\mu\text{M})$ |
| --- | --- | --- | --- | --- | --- |
| WT | $0.85 \pm 0.07$ | $0.37 \pm 0.09$ | $1.22 \pm 0.11$ | $1.15 \pm 0.13$ | $17.50 \pm 10.99$ |
| A92E | $0.92 \pm 0.08$ | $0.14 \pm 0.07$ | $1.06 \pm 0.11$ | $1.05 \pm 0.16$ | $5.75 \pm 12.17$ |
| $P$ vs. WT | 0.3206 | 0.1147 | 0.3618 | 0.4432 | 0.5133 |
| G562R | $0.10 \pm 0.05$ | $0.02 \pm 0.02$ | $0.12 \pm 0.05$ | $0.12 \pm 0.01$ | $4.10 \pm 21.35$ |
| $P$ vs. WT | 0.0001 | 0.0171 | 0.0008 | 0.0002 | 0.6066 |

Myo1e^A92E^ moved actin filaments 3-fold slower than Myo1e^WT^ (maximal unloaded sliding velocity of 109.7 ± 3.7 versus 297.1 ± 7.1 nm/s; Figure 5, Table 2). The Myo1e coating concentration required for half-maximal velocity was lower for Myo1e^A92E^ (0.10 ± 0.02 versus 0.17 ± 0.02 µM). Myo1e^G562R^ motility was not quantified due to its extremely low ATPase activity (Supplemental Movie 1-3).

**Figure 5.**
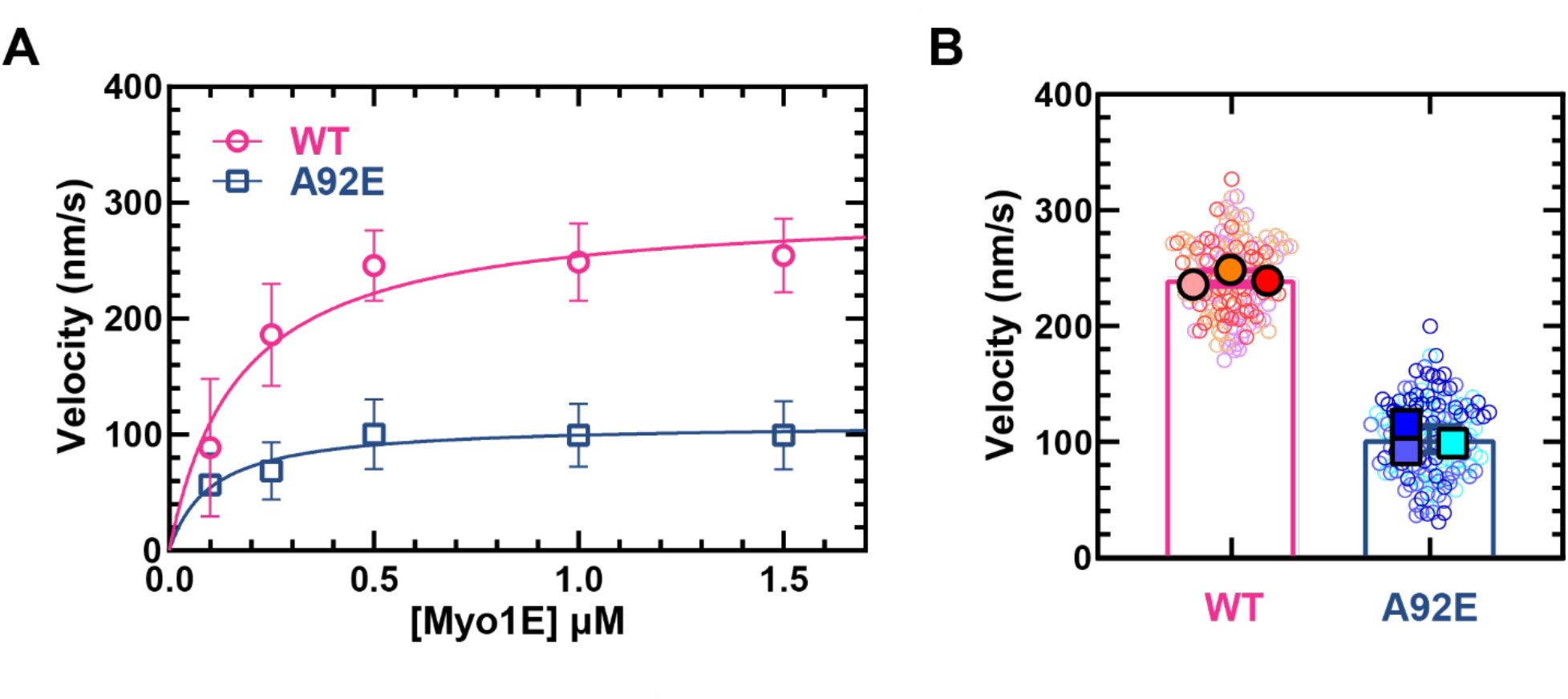
Myo1e-driven actin gliding velocity in vitro. (A) Actin filament gliding velocity as a function of myosin concentration for wild-type (WT; magenta circles) and A92E (blue squares) Myo1e. Solid lines represent hyperbolic fits to the data. Both WT and A92E exhibit increasing velocity with increasing myosin concentration, with A92E reaching a plateau at substantially lower velocities. Data points and error bars represent the mean ± standard deviation (SD) of velocities measured from 50 actin filaments at each myosin concentration. (B) Distribution of actin filament gliding velocities at a myosin concentration of 1 µM. Small open circles represent individual filament velocities, with different colors indicating three independent protein preparations. The filled symbols represent the mean velocity for each preparation. Bars and error bars indicate the mean ± SD across the three preparation means (N = 3; 50 filaments per preparation): WT, 241.6 ± 6.7 nm/s; A92E, 103.0 ± 11.5 nm/s.

**Table 2.** Summary of Myo1e in vitro motility measurements.

| Construct | Maximum gliding velocity<br>$V_{\max}$ (nm/s) | Density at which V is $\frac{1}{2} V_{\max}$<br>$K_{1/2}$ (μM) |
| --- | --- | --- |
| WT | 297.1 ± 7.1 | 0.17 ± 0.02 |
| A92E | 109.7 ± 3.7 | 0.10 ± 0.02 |
| *P* vs. WT | 0.0001 | 0.0150 |

### THERMOSTABILITY OF MYO1E DISEASE-ASSOCIATED VARIANTS

Thermostability assay of purified motor-IQ constructs showed a difference in the effects of the substitutions A92E and G562R on Myo1e stability (**Figure 6**, **Table 3**). The melting temperature of Myo1e^WT^ was 46.01 ± 0.19°C, while Myo1e A92E showed a substantially lower T_m_ of 34.31 ± 0.19°C. Thermal unfolding transition of Myo1e^G562R^ showed similar pattern to the WT with a T_m_ of 46.50 ± 0.22°C. These results indicate that the A92E mutation substantially reduces the thermal stability of Myo1e, whereas the G562R substitution has minimal effect on its global thermal stability.

**Figure 6.**
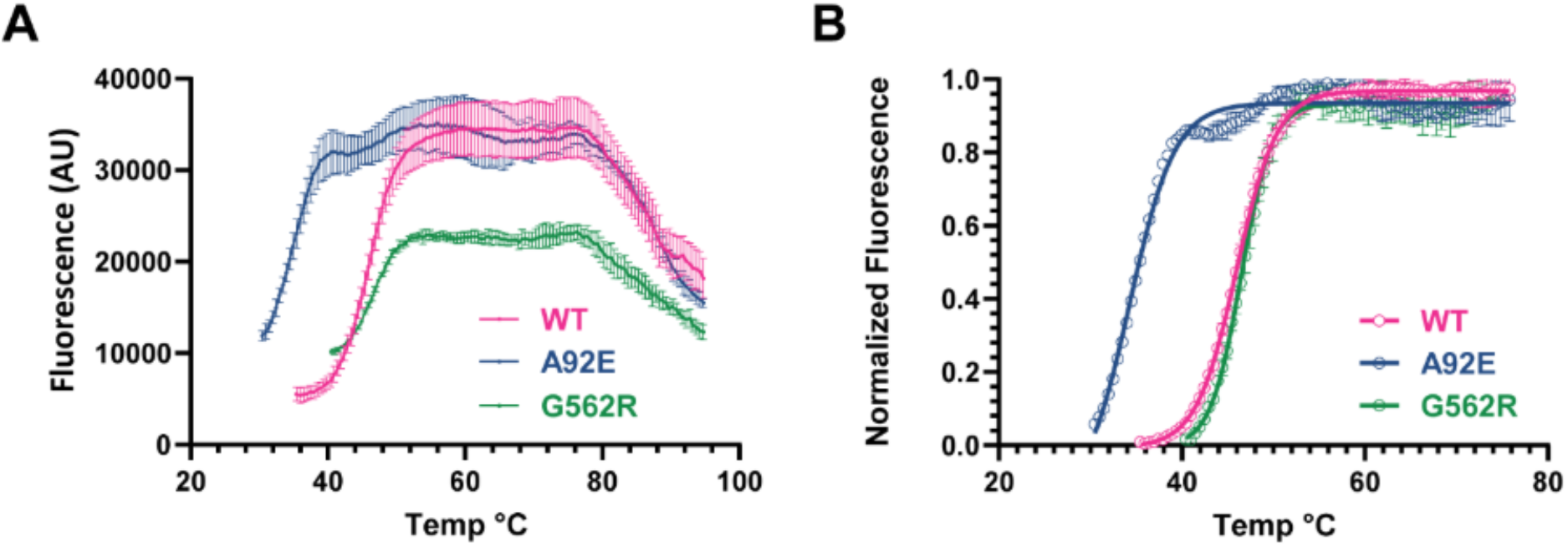
Thermal stability of Myo1e WT and disease-associated variants. (A) Raw fluorescence profiles of purified Myo1e WT, A92E, and G562R constructs measured using a dye-based thermal shift assay. (B) Normalized fluorescence curves showing the unfolding transitions. The normalized data were fitted using a Boltzmann sigmoidal equation, and the melting temperature (T_m_) was determined as the midpoint of the fitted transition. A92E exhibited a pronounced leftward shift relative to WT, whereas G562R showed a WT-like thermal stability profile. The calculated T_m_ values were 46.01 ± 0.19°C for WT, 34.31 ± 0.19°C for A92E, and 46.50 ± 0.22°C for G562R. Data are presented as mean ± SD from three technical replicates (n = 3).

**Table 3.**
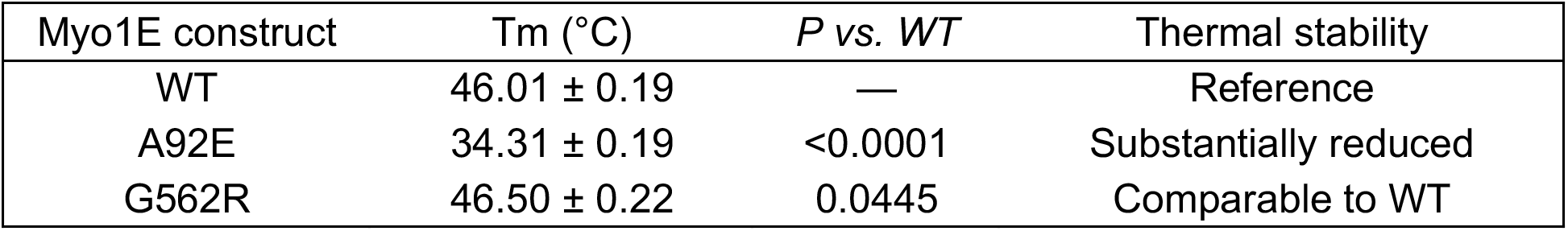
Thermal Stability of Myo1e WT and Disease-Associated Variants.

| Myo1E construct | $T_m$ ($^\circ\text{C}$) | $P$ vs. WT | Thermal stability |
| --- | --- | --- | --- |
| WT | $46.01 \pm 0.19$ | — | Reference |
| A92E | $34.31 \pm 0.19$ | $<0.0001$ | Substantially reduced |
| G562R | $46.50 \pm 0.22$ | 0.0445 | Comparable to WT |

## DISCUSSION

This study examined the effects of two disease-associated mutations in the motor domain of Myo1e, A92E and G562R, found in trans in a patient with childhood-onset SRNS and FSGS, on protein localization and motor activity, providing insight into how these variants may alter myosin function. Prior work (Liu et al., 2022) characterized two other Myo1e motor domain mutations associated with nephrotic syndrome, T119I and D388H. We found that the T119I substitution resulted in significant proteolytic degradation of the Myo1e motor-IQ fragment when expressed in insect cells and loss of localization in podocyte-derived cells while the D388H substitution impaired both ATPase activity and motility in vitro but did not interfere with the junctional localization of Myo1e in podocyte-like cells (Liu et al., 2022). Thus, it remains unclear whether FSGS-associated *MYO1E* variants converge on common functional defects, particularly with respect to enzymatic activity and subcellular localization. Our findings extend this analysis to additional clinically identified variants and suggest that disruption of Myo1e localization and motor properties may represent a recurring feature at both the biochemical and cellular levels.

Both Myo1e^A92E^ and Myo1e^G562R^ mutants failed to localize at cell-cell junctions, unlike wild-type Myo1e, which showed prominent enrichment at podocyte junctions and co-localization with cortical actin. Line-scan and junctional enrichment analyses confirmed a clear reduction in actin colocalization for both mutants, suggesting impaired membrane targeting or actin association. These findings are consistent with previous observations that Myo1e plays a critical role in maintaining podocyte junctional integrity and cytoskeletal architecture, and that its loss leads to podocyte effacement and proteinuria in knockout mice.

The Myo1e^G562R^ variant exhibited prominent cytoplasmic aggregation, distinct from Myo1e^A92E^ and WT. To determine whether these structures reflect formation of static protein aggregates, we performed FRAP on cytoplasmic Myo1e. Myo1e^G562R^ displayed a markedly higher immobile fraction and slower fluorescence recovery compared to the WT, consistent with reduced mobility within these aggregates. Lowering the expression level of Myo1e^G562R^ (by reducing viral vector concentration) partially suppressed aggregate formation without restoring junctional localization. This indicates that aggregation is partially dependent on the amount of mutant protein expressed in cells, but the failure to target junctions is likely an inherent property of the mutant protein, not just an overexpression artifact.

Using purified Myo1e constructs corresponding to the motor and IQ domains, we determined ATPase activity of the two SRNS-associated variants compared to that of the Myo1e^WT^. Myo1e^A92E^ retained most of the ATPase activity, while Myo1e^G562R^ had extremely low ATPase activity, indicating that this mutation severely affects ATPase activity. In the in vitro motility assay, the Myo1e^A92E^ mutant exhibited decreased ability to move actin filaments compared to the Myo1e^WT^. Thus, both mutations affect motor domain properties, such as the ability to hydrolyze ATP or move along actin filaments.

Both A92 and G562 are located within conserved α-helices of the myo1e motor domain. In silico modeling suggests that A92E and G562R may introduce local steric clashes and disrupt packing interactions critical for structural stability. While these predictions are consistent with the altered ATPase activity and motor behavior observed in vitro, they do not directly establish how these mutations perturb specific steps of the ATPase cycle or actin binding, which will require more detailed biophysical analysis. Our thermal stability measurements indicate that the A92E substitution also affects protein structural stability, which is consistent with the predictions above.

Given that Myo1e is enriched at the tips of the F-actin-rich podocyte foot processes, mutations that perturb its localization or interaction with actin could compromise the structural and signaling interface between podocytes and the glomerular basement membrane. The mis-localized and aggregation-prone mutants described here could compromise the dynamic coupling between the actin cytoskeleton and the plasma membrane at the slit diaphragm, a process thought to be essential for maintaining filtration barrier integrity under mechanical stress. Together with prior work on T119I and D388H, these results suggest that diverse *MYO1E* motor domain mutations converge on a limited set of functional defects, namely impaired junctional targeting and altered motor behavior, despite occurring at distinct structural positions.

In this study, we used a truncated Myo1e construct comprising the motor and neck domains, which enables controlled analysis of motor activity but does not capture the full regulatory and targeting functions of the tail. Full-length Myo1e is technically challenging to purify due to its size and conformational flexibility, and truncated constructs are therefore widely used in biochemical assays. However, future studies incorporating full-length protein will be important to determine how interdomain interactions and tail-dependent targeting contribute to Myo1e function in both normal and disease contexts. In addition, our intracellular localization analysis relies on overexpression in cultured cells, which may not fully recapitulate the stoichiometry and regulatory environment of endogenous Myo1e in podocytes, and future work may use physiologically relevant cell models such as gene-edited iPSC-derived podocytes.

Collectively, our findings suggest that disease-associated *MYO1E* mutations disrupt both motor function and subcellular targeting, supporting a model in which these coupled defects contribute to podocyte dysfunction in SRNS.

## ACKNOWLEDGMENTS

Molecular graphics and structural analyses were performed using UCSF ChimeraX. We thank Dr. Elaine Meng (UCSF ChimeraX) for expert guidance in protein modeling.

## GRANTS

This work was supported by the award R01DK083345 to MK.

## Author Contributions

KS - conceived and designed research, analyzed data, performed experiments, interpreted results of experiments, prepared figures, drafted manuscript, edited and revised manuscript, approved final version; JG - conceived and designed research, analyzed data, performed experiments, interpreted results of experiments, prepared figures, drafted manuscript, edited and revised manuscript, approved final version; CMY - conceived and designed research, analyzed data, interpreted results of experiments, drafted manuscript, edited and revised manuscript, approved final version; MK - conceived and designed research, interpreted results of experiments, prepared figures, drafted manuscript, edited and revised manuscript, approved final version.

---

**Supplemental Figure 1.**
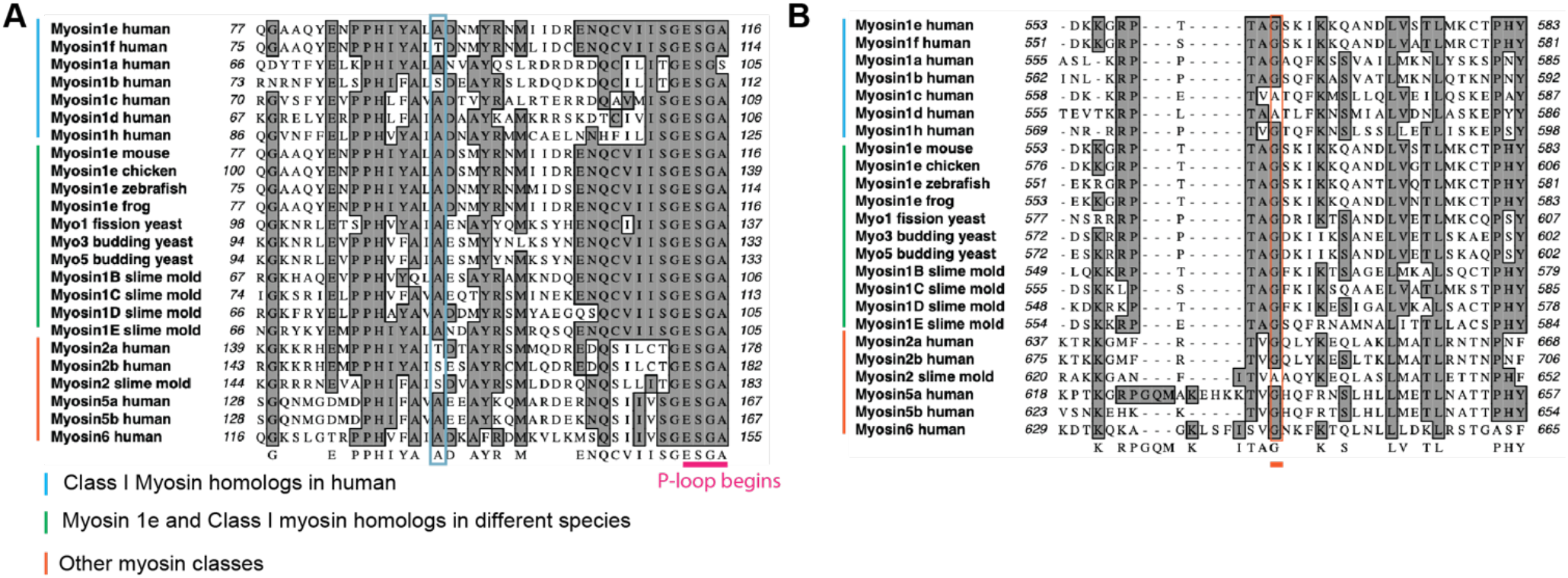
**Multiple sequence** alignment of regions surrounding A92 (A) and G562 (B) residues. The purple box in (**A**) and the orange box in (**B**) denote A92 and G562 residues, respectively. (**A**) A92 is highly conserved across Myo1e homologs in humans, class I myosins in different species, and other myosin classes, with substitutions with polar amino acids (serine or threonine) in human Myosin 2a and Myosin 2b, Dictyostelium discoideum (slime mold) Myosin 2, Myosin 1f and Myosin 1b in humans. (**B**) G562 is also broadly conserved across Myo1e homologs, class I myosins, and non-muscle myosin, except for Myosin2 in Dictyostelium discoideum (slime mold), Myosin 1c and Myosin 1d in humans, in which the corresponding position is occupied by a non-polar amino acid, alanine.

**Supplemental Table 1:**
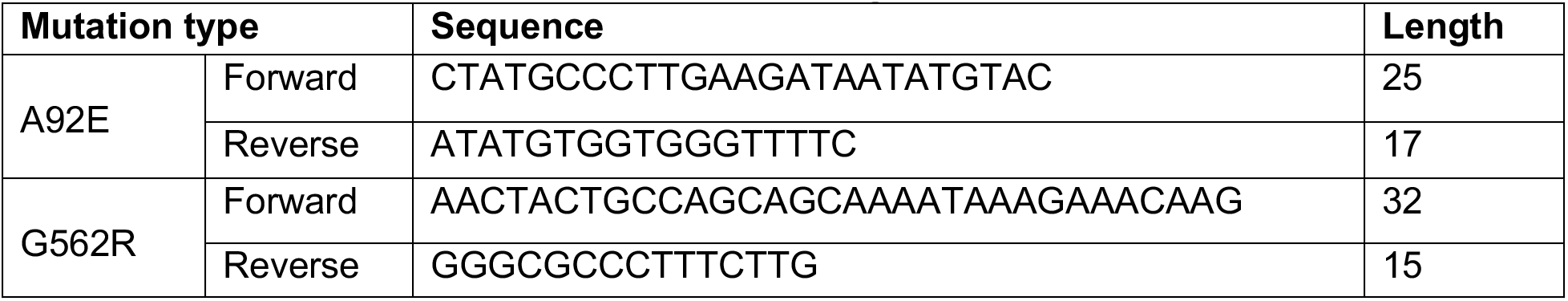
Primers used for mutagenesis.

**Supplemental Table 2:** NCBI reference sequence accession number.

| Protein | Species | Accession number |
| --- | --- | --- |
| Myo1e | Human (Homo sapiens) | NP_004989.2 |
| Myo1e | Mouse (Mus musculus) | NP_851417.2 |
| Myo1e | Zebrafish (Danio rerio) | NP_956930.1 |
| Myo1b | Human (Homo sapiens) | NP_036355.2 |
| Myo1f | Human (Homo sapiens) | CAC83948.1 |
| Myo1 | Fission yeast (Schizosaccharomyces pombe) | NP_595402.1 |
| Myo2a | Human (Homo sapiens) | NP_002464.1 |
| Myo1B | Slime mold (Dictyostelium discoideum) | XP_636382.1 |
| Myo5a | Human (Homo sapiens) | NP_000250.3 |
| Myo6 | Human (Homo sapiens) | NP_004990.3 |
| Myo1a | Human (Homo sapiens) | NP_001242970.1 |
| Myo1b | Human (Homo sapiens) | NP_036355.2 |
| Myo1c | Human (Homo sapiens) | NP_203693.3 |
| Myo1d | Human (Homo sapiens) | NP_056009.1 |
| Myo1h | Human (Homo sapiens) | NP_001094891.4 |
| Myo1e | Chicken (Gallus gallus) | XP_413782.4 |
| Myo1e | African frog (Xenopus laevis) | XP_018108508.1 |
| Myo3 | Budding yeast (Saccharomyces cerevisiae) | NP_012793.2 |
| Myo5 | Budding yeast (Saccharomyces cerevisiae) | NP_013827.1 |
| Myo1C | Slime mold (Dictyostelium discoideum) | XP_643060.1 |
| Myo1D | Slime mold (Dictyostelium discoideum) | XP_643446.1 |
| Myo1E | Slime mold (Dictyostelium discoideum) | XP_636580.1 |
| Myo2b | Human (Homo sapiens) | NP_001242941.1 |
| Myo2 | Slime mold (Dictyostelium discoideum) | XP_637740.1 |
| Myo5b | Human (Homo sapiens) | NP_001073936.1 |

**Supplemental Movie 1.** In vitro motility of wild-type (WT) Myo1e (1 μM) in motility buffer. Time-lapse images were acquired at 10-second intervals for up to 10 minutes.

**Supplemental Movie 2.** In vitro motility of the Myo1e^A92E^ mutant under the same experimental and imaging conditions as Supplemental Movie 1.

**Supplemental Movie 3.** In vitro motility of the Myo1e^G562R^ mutant under the same experimental and imaging conditions as Supplemental Movie 1.

## Notes

### Competing Interest Statement

The authors have declared no competing interest.

## REFERENCES

Ali, S.A., M.I. Hassan, A. Islam, and F. Ahmad. 2014. A review of methods available to estimate solvent-accessible surface areas of soluble proteins in the folded and unfolded states. Curr Protein Pept Sci. 15:456–476.

Aloy, P., E. Querol, F.X. Aviles, and M.J. Sternberg. 2001. Automated structure-based prediction of functional sites in proteins: applications to assessing the validity of inheriting protein function from homology in genome annotation and to protein docking. J Mol Biol. 311:395–408.

Bi, J., S.E. Chase, C.D. Pellenz, H. Kurihara, A.S. Fanning, and M. Krendel. 2013. Myosin 1e is a component of the glomerular slit diaphragm complex that regulates actin reorganization during cell-cell contact formation in podocytes. Am J Physiol Renal Physiol. 305:F532–544.

Bi, J., C.D. Pellenz, and M. Krendel. 2014. Visualization of cytoskeletal dynamics in podocytes using adenoviral vectors. Cytoskeleton (Hoboken). 71:145–156.

Bierzynska, A., and M. Saleem. 2017. Recent advances in understanding and treating nephrotic syndrome. F1000Res. 6:121.

Cason, R.K., E. Chambers, T. Tu, M. Chryst-Stangl, K. Huggins, B.M. Lane, A. Ochoa, A.M. Jackson, and R.A. Gbadegesin. 2023. Genetic risk variants for childhood nephrotic syndrome and corticosteroid response. Front Pediatr. 11:1248733.

Chase, S.E., C.V. Encina, L.R. Stolzenburg, A.H. Tatum, L.B. Holzman, and M. Krendel. 2012. Podocyte-specific knockout of myosin 1e disrupts glomerular filtration. Am J Physiol Renal Physiol. 303:F1099–1106.

Dantas, V.G.L., M.H. Raval, A. Ballesteros, R. Cui, L.K. Gunther, G.L. Yamamoto, L.U. Alves, A.S. Bueno, K. Lezirovitz, S. Pirana, B.C.A. Mendes, C.M. Yengo, B. Kachar, and R.C. Mingroni-Netto. 2018. Characterization of a novel MYO3A missense mutation associated with a dominant form of late onset hearing loss. Sci Rep. 8:8706.

De La Cruz, E.M., H.L. Sweeney, and E.M. Ostap. 2000. ADP inhibition of myosin V ATPase activity. Biophys J. 79:1524–1529.

Diensthuber, R.P., F.K. Hartmann, D. Kathmann, P. Franz, and G. Tsiavaliaris. 2024. Switch-2 determines Mg(2+)ADP-release kinetics and fine-tunes the duty ratio of Dictyostelium class-1 myosins. Front Physiol. 15:1393952.

Eddy, A.A., and J.M. Symons. 2003. Nephrotic syndrome in childhood. Lancet. 362:629–639.

El Mezgueldi, M., N. Tang, S.S. Rosenfeld, and E.M. Ostap. 2002. The kinetic mechanism of Myo1e (human myosin-IC). J Biol Chem. 277:21514–21521.

Gress, A., and O.V. Kalinina. 2020. SphereCon-a method for precise estimation of residue relative solvent accessible area from limited structural information. Bioinformatics. 36:3372–3378.

Gupta, P., N.C. Gauthier, Y. Cheng-Han, Y. Zuanning, B. Pontes, M. Ohmstede, R. Martin, H.J. Knolker, H.G. Dobereiner, M. Krendel, and M. Sheetz. 2013. Myosin 1E localizes to actin polymerization sites in lamellipodia, affecting actin dynamics and adhesion formation. Biology open. 2:1288–1299.

Ha, T.S. 2017. Genetics of hereditary nephrotic syndrome: a clinical review. Korean J Pediatr. 60:55–63.

Hall, G., and R.A. Gbadegesin. 2015. Translating genetic findings in hereditary nephrotic syndrome: the missing loops. Am J Physiol Renal Physiol. 309:F24–28.

Heim, J.B., E.J. Squirewell, A. Neu, G. Zocher, S. Sominidi-Damodaran, S.P. Wyles, E. Nikolova, N. Behrendt, D.M. Saunte, J. Lock-Andersen, K.S. Gaonkar, H. Yan, J.N. Sarkaria, M. Krendel, J. van Deursen, R. Sprangers, T. Stehle, R.T. Bottcher, J.H. Lee, T. Ordog, and A. Meves. 2017. Myosin-1E interacts with FAK proline-rich region 1 to induce fibronectin-type matrix. Proc Natl Acad Sci U S A. 114:3933–3938.

Heiringhoff, R.S., J.N. Greve, M. Zahn, and D.J. Manstein. 2025. Structure of the human nonmuscle myosin 2A motor domain: Insights into isoform-specific mechanochemistry. J Biol Chem. 301:110691.

Kintses, B., M. Gyimesi, D.S. Pearson, M.A. Geeves, W. Zeng, C.R. Bagshaw, and A. Malnasi-Csizmadia. 2007. Reversible movement of switch 1 loop of myosin determines actin interaction. EMBO J. 26:265–274.

Krendel, M., S.V. Kim, T. Willinger, T. Wang, M. Kashgarian, R.A. Flavell, and M.S. Mooseker. 2009. Disruption of Myosin 1e promotes podocyte injury. Journal of the American Society of Nephrology : JASN. 20:86–94.

Krendel, M., S. Leh, M.E. Garone, A. Edwards-Richards, J.J. Lin, D. Brackman, P. Knappskog, and A. Mikhailov. 2023. Focal segmental glomerulosclerosis and proteinuria associated with Myo1E mutations: novel variants and histological phenotype analysis. Pediatr Nephrol. 38:439–449.

Krendel, M., E.K. Osterweil, and M.S. Mooseker. 2007. Myosin 1E interacts with synaptojanin-1 and dynamin and is involved in endocytosis. FEBS Lett. 581:644–650.

Liu, P.J., L.K. Gunther, M.E. Garone, C. Zhang, D. Perez, J. Bi-Karchin, C.D. Pellenz, S.E. Chase, M.F. Presti, E.L. Plante, C.E. Martin, S. Lovric, C.M. Yengo, F. Hildebrandt, and M. Krendel. 2022. Steroid-Resistant Nephrotic Syndrome-Associated MYO1E Mutations Have Differential Effects on Myosin 1e Localization, Dynamics, and Activity. Journal of the American Society of Nephrology : JASN. 33:1989–2007.

Liu, P.J., K. Sayeeda, C. Zhuang, and M. Krendel. 2024. Roles of myosin 1e and the actin cytoskeleton in kidney functions and familial kidney disease. Cytoskeleton (Hoboken). 81:737–752.

McIntosh, B.B., and E.M. Ostap. 2016. Myosin-I molecular motors at a glance. J Cell Sci. 129:2689–2695.

Meijering, E., O. Dzyubachyk, and I. Smal. 2012. Methods for cell and particle tracking. Methods Enzymol. 504:183–200.

Mele, C., P. Iatropoulos, R. Donadelli, A. Calabria, R. Maranta, P. Cassis, S. Buelli, S. Tomasoni, R. Piras, M. Krendel, S. Bettoni, M. Morigi, M. Delledonne, C. Pecoraro, I. Abbate, M.R. Capobianchi, F. Hildebrandt, E. Otto, F. Schaefer, F. Macciardi, F. Ozaltin, S. Emre, T. Ibsirlioglu, A. Benigni, G. Remuzzi, M. Noris, and C. PodoNet. 2011. MYO1E mutations and childhood familial focal segmental glomerulosclerosis. The New England journal of medicine. 365:295–306.

Meng, E.C., T.D. Goddard, E.F. Pettersen, G.S. Couch, Z.J. Pearson, J.H. Morris, and T.E. Ferrin. 2023. UCSF ChimeraX: Tools for structure building and analysis. Protein Sci. 32:e4792.

Mercier, E., D. Girodat, and H.J. Wieden. 2015. A conserved P-loop anchor limits the structural dynamics that mediate nucleotide dissociation in EF-Tu. Sci Rep. 5:7677.

Mirdita, M., K. Schutze, Y. Moriwaki, L. Heo, S. Ovchinnikov, and M. Steinegger. 2022. ColabFold: making protein folding accessible to all. Nat Methods. 19:679–682.

Mundel, P., J. Reiser, A. Zuniga Mejia Borja, H. Pavenstadt, G.R. Davidson, W. Kriz, and R. Zeller. 1997. Rearrangements of the cytoskeleton and cell contacts induce process formation during differentiation of conditionally immortalized mouse podocyte cell lines. Exp Cell Res. 236:248–258.

Noris, M., and G. Remuzzi. 2012. Non-muscle myosins and the podocyte. Clin Kidney J. 5:94–101.

Pardee, J.D., and J.A. Spudich. 1982. Purification of muscle actin. Methods Enzymol. 85 Pt B:164–181.

Pettersen, E.F., T.D. Goddard, C.C. Huang, E.C. Meng, G.S. Couch, T.I. Croll, J.H. Morris, and T.E. Ferrin. 2021. UCSF ChimeraX: Structure visualization for researchers, educators, and developers. Protein Sci. 30:70–82.

Preston, R., H.M. Stuart, and R. Lennon. 2019. Genetic testing in steroid-resistant nephrotic syndrome: why, who, when and how? Pediatr Nephrol. 34:195–210.

Raval, M.H., O.A. Quintero, M.L. Weck, W.C. Unrath, J.W. Gallagher, R. Cui, B. Kachar, M.J. Tyska, and C.M. Yengo. 2016. Impact of the Motor and Tail Domains of Class III Myosins on Regulating the Formation and Elongation of Actin Protrusions. J Biol Chem. 291:22781–22792.

Sanna-Cherchi, S., K.E. Burgess, S.N. Nees, G. Caridi, P.L. Weng, M. Dagnino, M. Bodria, A. Carrea, M.A. Allegretta, H.R. Kim, B.J. Perry, M. Gigante, L.N. Clark, S. Kisselev, D. Cusi, L. Gesualdo, L. Allegri, F. Scolari, V. D’Agati, L.S. Shapiro, C. Pecoraro, T. Palomero, G.M. Ghiggeri, and A.G. Gharavi. 2011. Exome sequencing identified MYO1E and NEIL1 as candidate genes for human autosomal recessive steroid-resistant nephrotic syndrome. Kidney Int. 80:389–396.

Savojardo, C., M. Manfredi, P.L. Martelli, and R. Casadio. 2020. Solvent Accessibility of Residues Undergoing Pathogenic Variations in Humans: From Protein Structures to Protein Sequences. Front Mol Biosci. 7:626363.

Sun, M., J.L. Oakes, S.K. Ananthanarayanan, K.H. Hawley, R.Y. Tsien, S.R. Adams, and C.M. Yengo. 2006. Dynamics of the upper 50-kDa domain of myosin V examined with fluorescence resonance energy transfer. J Biol Chem. 281:5711–5717.

Tang, W., J. Ge, W.C. Unrath, R. Desetty, and C.M. Yengo. 2021. Cardiomyopathy mutations impact the actin-activated power stroke of human cardiac myosin. Biophys J. 120:2222–2236.

Tien, M.Z., A.G. Meyer, D.K. Sydykova, S.J. Spielman, and C.O. Wilke. 2013. Maximum allowed solvent accessibilites of residues in proteins. PLoS One. 8:e80635.

Trautmann, A., M. Vivarelli, S. Samuel, D. Gipson, A. Sinha, F. Schaefer, N.K. Hui, O. Boyer, M.A. Saleem, L. Feltran, J. Muller-Deile, J.U. Becker, F. Cano, H. Xu, Y.N. Lim, W. Smoyer, I. Anochie, K. Nakanishi, E. Hodson, D. Haffner, and A. International Pediatric Nephrology. 2020. IPNA clinical practice recommendations for the diagnosis and management of children with steroid-resistant nephrotic syndrome. Pediatr Nephrol. 35:1529–1561.

Trivedi, D.V., J.M. Muretta, A.M. Swenson, D.D. Thomas, and C.M. Yengo. 2013. Magnesium impacts myosin V motor activity by altering key conformational changes in the mechanochemical cycle. Biochemistry. 52:4710–4722.

van der Velde, K.J., H.S. Dhekne, M.A. Swertz, S. Sirigu, V. Ropars, P.C. Vinke, T. Rengaw, P.C. van den Akker, E.H. Rings, A. Houdusse, and S.C. van Ijzendoorn. 2013. An overview and online registry of microvillus inclusion disease patients and their MYO5B mutations. Hum Mutat. 34:1597–1605.

Varghese, R., and A. Majumdar. 2022. A New Prospect for the Treatment of Nephrotic Syndrome Based on Network Pharmacology Analysis. Curr Res Physiol. 5:36–47.

Xiong, D., D. Lee, L. Li, Q. Zhao, and H. Yu. 2022. Implications of disease-related mutations at protein-protein interfaces. Curr Opin Struct Biol. 72:219–225.

Yengo, C.M., Y. Takagi, and J.R. Sellers. 2012. Temperature dependent measurements reveal similarities between muscle and non-muscle myosin motility. J Muscle Res Cell Motil. 33:385–394.

